# Glycation of alpha-synuclein enhances aggregation and neuroinflammatory responses

**DOI:** 10.1101/2024.06.27.600956

**Authors:** Eftychia Vasili, Annekatrin König, Mohammed Al-Azzani, Clara Bosbach, Luisa Maria Gatzemeier, Ana Chegão, Hugo Vicente Miranda, Daniel Erskine, Tiago F. Outeiro

**Affiliations:** University Medical Center Gottingen, Department of Experimental Neurodegeneration, Center for Biostructural Imaging of Neurodegeneration, Göttingen, Germany; Institute of Organic and Biomolecular Chemistry, Georg-August-University Göttingen, Tammannstraße 2, 37077 Göttingen, Germany; iNOVA4Health, NOVA Medical School, NMS, Universidade NOVA de Lisboa, Lisboa, Portugal; Translational and Clinical Research Institute, Newcastle University, Newcastle upon Tyne NE1 7RU, United Kingdom; Max Planck Institute for Multidisciplinary Sciences, Göttingen, Germany; Scientific employee with an honorary contract at Deutsches Zentrum für Neurodegenerative Erkrankungen (DZNE), Göttingen, Germany

**Keywords:** Parkinson’s disease, alpha-synuclein, neurodegeneration, glycation, protein aggregation, diabetes

## Abstract

The risk of developing Parkinson’s disease (PD) is elevated in people with type 2 diabetes, but the precise molecular pathways underlying this connection are still unclear. One hypothesis is that glycation, a non-enzymatic family of reactions between glycating agents, such as reducing sugars or reactive dicarbonyls, and specific amino acids, such as lysines and arginines, may alter proteostasis and trigger pathological alterations. Glycation of alpha-synuclein (aSyn), a central player in PD pathology, causes profound changes in the aggregation process of aSyn. Methylglyoxal (MGO), a strong glycating agent, induces the formation of pathological inclusions enriched in phosphorylated aSyn on serine 129 (pS129). In addition, we found that neuroinflammatory responses are enhanced by MGO-mediated aSyn glycation. Using novel polyclonal antibodies developed towards specific MGO-glycated aSyn residues, we confirmed the occurrence of glycated aSyn both *in vitro* as well as in animal and in human brain tissue. In total, our findings shed light into the interplay between glycation, PD, and type 2 diabetes, potentially paving the way for the development of novel therapeutic strategies targeting these intertwined conditions.

## Introduction

Glycation, also referred to as non-enzymatic glycosylation or Maillard reaction, is the reaction between reducing sugars or other aldehydes and amino groups or nucleotides ^1^. Glycation is actually a complex network of different reactions, that involve the formation of Schiff bases in the first steps ^2^. Early glycation products, also called Amadori compounds, are formed via condensation of a carbonyl form with an amino or thiol group of amino acids, nucleic acids or amino lipids. The intermediate Amadori product is then rearranged and converted in one of different chemical pathways^3^. Several different intermediary products, including glyoxal and methylglyoxal (MGO), can be formed. Finally, higher and irreversible molecular weight species (advanced glycation end products – AGEs) are generated^4^. Proteins and peptides can be modified at the *N*-terminus or at arginine or lysine side chains. However, since glycation is a non-enzymatic reaction, it is not limited to defined residues or to physiological situations. Although several reducing sugars, such as glucose, fructose, and ribose, can contribute to glycation processes, the most efficient glycation agent *in vivo* is MGO, that is formed as a side product in glycolysis ^5–7^ and from the catabolism of amino acids.

Alpha-Synuclein (aSyn), a central player in the etiology and progression of Parkinsońs disease (PD) and related disorders (collectively known as synucleinopathies) is, like many other proteins, subject to glycation reactions ^8^ However, given that aSyn is rich in lysines (15 residues in its primary amino acid sequence), and particularly long-lived, aSyn-AGEs can form and accumulate ^9^. We and others have previously shown that glycation affects several aspects of aSyn biology. In particular, glycation impairs aSyn clearance due to interference with ubiquitination and SUMOylation of lysine residues, reduces extracellular protease cleavage, and impairs membrane binding ^10–12^. MGO-mediated glycation was shown to affect mainly the aSyn *N*-terminal lysine residues ^10^. Interestingly, glycation affects aSyn aggregation kinetics, enhancing oligomerization and reducing fibrilization ^8,10,13–16^. Around 5 MGO-glycated residues per aSyn molecule were shown to optimally inhibit aSyn aggregate elongation, reducing the incorporation of further aSyn monomers into the fibrils ^8^. Strikingly, intracerebroventricular injection of MGO aggravates motor, cognitive, olfactory, and colonic dysfunction in a mouse model of synucleinopathy ^17^.

The metabolic disease diabetes mellitus type 2 (T2DM), characterized by sugar dyshomeostasis and increased levels of protein glycation, is a known risk factor for PD ^18–21^. Given the particular vulnerability of neurons to high glucose levels, glycation, and associated oxidative stress, understanding the effects of glycation in proteins of interest in the context of brain disorders is essential. Apart from the direct consequences of hyperglycemia that are thought to account for most of the T2DM associated complications, higher glucose levels also cause increased production of reactive oxoaldehydes such as MGO ^6,22^. In addition, abnormally high levels of D-ribose, an important contributor to protein glycation, are detected in the urine of T2DM patients ^23^. Thus, in this study we focused on the molecular effects of these two agents (MGO and ribose) on aSyn spreading, seeding and neuroinflammation. In addition, we developed novel aSyn glycation-specific polyclonal antibodies that will boost our understanding of the effects of aSyn glycation in PD and in other synucleinopathies. Ultimately, we anticipate that our findings may open novel perspectives for therapeutic intervention.

## Materials and Methods

### Production and purification of aSyn

Production and purification of recombinant aSyn was performed as described ^24^. In brief, pET21-aSyn was transformed into competent *E. coli* BL21-DE3 (Sigma), *E. coli* was grown in 2x LB medium with ampicillin (200 μg/mL) at 37°C with constant shaking. At an OD_600_ of 0.5–0.6, the expression was induced with 1 mM of isopropylβ-thiogalactopyranoside. After 2 hours, bacteria were pelleted by centrifugation at 6600g, 15 min and lysed on ice in 10 mM Tris pH7.6, 750 mM NaCl, 1 mM EDTA with protease inhibitor (cOmplete, Roche). Lysates were sonicated on ice for a total of 5 min (30s on, 30s off pulses, 60% power), and heated for 15 min at 95 °C. Samples were then centrifuged at 15.000g and the supernatant was subjected to dialysis in 10 mM TRIS pH 7.6, 1 mM EDTA, 50 mM NaCl. Anion exchange chromatography (HiTrap Q HP, GE Healthcare) was performed with a mobile phase of 25 mM Tris pH 7.6 and a linear gradient of 9 column volumes of elution buffer to 1M NaCl on an Äkta Pure 25M (Cytiva). Fractions containing pure aSyn were identified on a Coomassie-stained SDS-PAGE and pooled. Proteins were further purified with size exclusion chromatography using a hiLoad Superdex200pg column (Cytiva). Fractions containing pure aSyn were identified on a Coomassie-stained SDS-PAGE and pooled and extensively dialyzed into water. Following lyophilization (Zirbus), protein was stored at −20°C. aSyn concentration was measured using absorbance at 280nm (molar extinction coefficient 5960 M^−1^ cm^−1^).

### Glycation of recombinant aSyn

100mM aSyn was dissolved in 1xPBS, 7.4 with 3.7 mM EDTA with or without 5mM MGO (Sigma) or 0.8mM ribose and filtered with 0.2 μM membrane syringe filter. The samples were then incubated for five days at 37 °C, under constant agitation (300 rpm) in low binding tubes (Corning Incorporated). Glycation was confirmed via measuring fluorescence as described in ^8^. Briefly, Fluorescence of glycation products was measured by excitation at 340 nm and emission at 390 nm.

### Cell cultures

The human neuroblastoma (SH-SY5Y) cells used conditionally expressed human wildtype aSyn under a Tet-Off cassette ^25^. Cells were grown in RPMI with 1 μM doxycycline, 10% FBS, and 1% penicillin/streptomycin at 37 °C, 5% CO2. To induce aSyn expression from the Tet-Off cassette, doxycycline was omitted. For differentiation, cells were seeded in 6-well plates. One day after plating, the medium was supplemented with 10 μM retinoic acid in RPMI supplemented with 0.5% FBS and 1% penicillin/streptomycin, at 37 °C, 5% CO2 for 4 days. Media was changed every second day. Following differentiation, cells were treated with 100 nM of the respective glycated aSyn species for 4 days. aSyn treated the same way but without glycating agents was used as control. Mild trypsinization for 5 min was used to remove excess unbound aSyn protein. Cultures were washed with PBS and fixed with 4% paraformaldehyde (PFA) for 20 min at RT.

### Primary rat neuronal cultures

7 to 12 primary cortical neurons were prepared from embryonic day E18 Wistar rat brains. After dissection in Hank’s Balanced Salt Solution (HBSS; Gibco), the cortices were digested by Trypsin (Gibco) for 15 min at 37 °C, followed by addition of 100μl DNase I (10 mg/ml; Roche) and 100μl FBS (Life Technologies), mixed by inverting and spun down. The liquid was replaced with 1ml FBS and the tissue triturated with a pasteur pipette. The dissociated cells were spun down and transferred into fresh culture medium (Neurobasal medium, Gibco), with 2% B27 supplement (Gibco), 0.5mM L-glutamine (200Mm Gibco), and 1% penicillin/streptomycin (Gibco). Cells were plated in in 24-or 6-well plates previously coated with Poly-L-ornithin (0,1 mg/ml, Sigma-Aldrich) at a density of 250,000 cells/mL. The cultures were differentiated for 5 days. At day 5, aSyn species were added to a final concentration of 50nM and incubated for 20 days. Mild trypsinization for 5 min was used to remove excess unbound aSyn material. Cultures were washed with PBS and fixed with 4% paraformaldehyde (PFA) for 20 min at RT.

### Primary microglial cultures

Primary microglia were obtained from mixed glial cell cultures from C57BL6 wild-type newborn mice as described previously ^26^. Briefly, brains were isolated in 1xHBSS (Gibco Invitrogen), meninges were removed and the brains were washed three times with HBSS (without Ca2 +, Mg2 + and phenol; PAN Biotech). The tissue was trypsinized (0.05% trypsin-EDTA; PAN Biotech) at 37°C for 10 minutes. After aspirating the trypsin solution, the digestion was stopped by adding 0.5 mg/mL DNase I (Roche) in microglia medium (DMEM (PAN Biotech), supplemented with 0.5% penicillin-streptomycin (PAN Biotech) and 10% FBS (Anprotec)). Tissue was incubated for three further minutes at 37°C and homogenized with a glass pipette. Cells were spun down at 800g for 10 minutes, the pellet was resuspended in medium. Cells were plated into T75 flasks (Corning, Merck) that had previously been coated with poly-L-ornithine (Sigma-Aldrich, 0.1 mg/mL in borate buffer) overnight at 37°C and 5% CO2. The following day (DIV2), the cells were washed three times with pre-warmed HBSS solution (PAN Biotech) and once with microglia medium before new medium was added. At DIV3, cell medium was replaced once more. At DIV5, the culture was stimulated by substituting one third of the medium with L929 medium (L929 mouse fibroblast cells had previously been plated in T175 cm2 cell culture flasks (Corning, Merck) with 100 mL culture medium (DMEM, PAN biotech; supplemented with 10% FBS (Anprotec) and 1% penicillin-streptomycin (PAN Biotech)) for seven days at 37°C with 5% CO2. The media was then collected, sterilized by filtration using a 0.22μm filter (Sartorius, Göttingen, Germany) and stored at –20°C). At DIV8, microglia were harvested by mild shaking and collected (10 minutes at 800g) and replated into culture plates previously coated with poly-L-ornithine (Sigma-Aldrich, 0.1 mg/mL in borate buffer). Microglia were allowed to attach to the new flasks for 16 hours and treated with 100nM aSyn species for 24 hours.

### Production of polyclonal antibodies

The sequences of peptides 1 (MKGLS KAKEG VVAAA EKTK) and 2 (KTKEG VLYVG SKTKE GVVHG VATVA EKTK) were determined in cooperation with experts from Davids Biotechnology based on lysine residues in aSyn. Full-length aSyn, peptide 1 or peptide 2 were glycated as described above and were used to immunize rabbits by Davids Biotechnology, Regensburg, according to established immunization plans. Selected antibodies were then affinity purified.

### RNA isolation and qRT-PCR

RNA was extracted from microglial cultures using TRIzol Reagent according to the manufacturer’s instructions (Invitrogen). cDNA was reverse transcribed using QuantiTect Reverse Transcription kit (Qiagen, MD, USA) according to the manufacturer’s instructions. qPRC was performed on an Applied Biosystems Real-Time PCR Systems using SYBR Green Master Mix (Qiagen); 95°C for 10 min, then 40 cycles at 95°C for 15 s and 60°C for 25 s. The following primers were used: IL-6 F: ATCCA GTTGC CTTCT TGGGA CTGA, IL-6 R:TAAGC CTCCG ACTTG TGAAG TGGT, IL-1β F: TCATT GTGGC TGTGG AGAAG, IL-1β R: AGGCC ACAGG TATTT TGTCG, TNFα F: CCCTC TCATC AGTTC TATGG, TNFα R: GGAGT AGACA AGGTA CAACC, *β*-actin F: GCGAG AAGAT GACCC AGATC, and *β*-actin R: CCAGT GGTAC GGCCA GAGG. *β*-actin was used as reference gene to calculate the fold change in expression levels using the 2–ΔΔCT method.

### Dot blots

For the dot blots, the protein/peptide specified was diluted in 100µL PBS and spotted onto a nitrocellulose membrane, (0.1µm for peptides, 0.2µm for proteins; Amersham Protran) using a custom-made dot blot apparatus with vacuum. The membranes were completely air-dried and incubated at room temperature with blocking solution. If peptides were used, performing aSyn staining as a loading control was not possible. In this case, the Pierce™ Reversible Protein Stain Kit was used according to manufacturer’s instructions.

Primary antibodies were incubated over night at 4°C with mild agitation. The polyclonal antibodies generated were used in a concentration of 1µg/mL and diluted in blocking buffer (0.1M sodium acetate, 5% BSA, 0.05% tween 20). Syn1 (BD Transduction Laboratories, 1:1000) was used to assess total aSyn concentration. TBS-Tween (1 × TBS, supplemented with 0.05% (v/v) Tween-20) was used for three 15-minute washes. Secondary antibodies (anti-mouse and anti-rabbit IgG, 1∶10,000 in blocking buffer) were applied for one hour at room temperature. After three more washes in TBS-Tween, membranes were developed using Fusion Fx (Vilber Lourmat) with Immobilon Western Chemiluminescent HRP Substrate (Merck Millipore).

### Sequential protein extraction

Cell pellets were collected and solubilized in 1% Triton X-buffer (50 mM tris pH 7.6, 150 mM NaCl, 2 mM EDTA, 1% Triton X-100, protease-and phosphatase inhibitors), followed by a 30 minute incubation on ice. Cell lysates were spun down at 13,000 g for 30 minutes, 4 °C. The supernatant is the Triton X soluble fraction. The pellet was washed with ice cold PBS, resuspended in 2% SDS buffer (50 mM tris pH 7.6, 150 mM NaCl, 2 mM EDTA, 2% SDS, protease-and phosphatase inhibitors), sonicated and incubated at room temperature for 30 minutes. After centrifugation at 13,000 g for 10 min, the supernatant - the SDS soluble fraction – was collected. Bradford protein assay was used to determine the protein concentration and the samples were then subjected to Western blot analysis.

### SDS-PAGE and immunoblotting

For total protein cell lysates, cells were washed with PBS and lysed on ice in radio-immunoprecipitation assay buffer (RIPA) (50 mM Tris pH 8.0, 150 mM NaCl, 0.1% Sodium-Dodecyl-Sulphate (SDS), 1% Nonidet P40, 0.5% Sodium-Deoxycholate, protease inhibitors, Roche Diagnostics, Mannheim, Germany). Lysates were centrifuged at 10000rpm and 4 °C for 10 min and post-nuclear supernatants were kept. Protein concentration was determined using the Bradford assay (BioRad). All samples were measured in triplicate. Equal protein amounts of denatured samples (5 minutes at 95°C in 5x protein sample buffer; 125 mM of 1 M Tris HCl pH 6.8, 4% SDS 0,5% Bromophenol blue, 4 mM EDTA 20% Glycerol 10% β-Mercaptoethanol) were subjected to SDS-PAGE on 12% separating gels with 7% stacking gels, using Tris-Glycine SDS 0.5% running buffer (250 mM Tris, 200 mM Glycine, 1% SDS, pH 8.3). The transfer was carried out to 0.45 μm nitrocellulose membranes for 20 min per membrane at constant 25 mA in a semi-dry transfer chamber Trans-Blot® Turbo™ Transfer Solution from Bio-Rad (Bio-Rad Laboratories, Inc., Hercules, CA, USA). Membranes were blocked in 5% (w/v) skim milk (Fluka, Sigma-Aldrich, St. Louis, MO, USA) dissolved in 1xTBS-Tween (50 mM Tris (hydroxymethyl)-aminomethane (TRIS) supplemented with 0.05% (v/v) Tween-20) for 1 h at RT. Incubation with the primary antibodies (anti-aSyn Syn-1 mouse, BD Transduction Laboratories 1:1000; anti-pS129-α-syn Rabbit Abcam Ab51253; mouse anti-*β*-actin, 1:10.000, Sigma Aldrich) was performed overnight at 4 °C in 5% Albumin Bovine Fraction V (BSA)/TBS-Tween. Secondary antibodies (anti-mouse and anti-rabbit IgG, 1∶10000 in TBS-Tween) were applied after three times washing in TBS-Tween, for 1 h at RT. Membranes were visualized using Fusion Fx (Vilber Lourmat, Marne-la-Vallée, France) with Immobilon Western Chemiluminescent HRP Substrate (Merck Millipore, Billerica, MA, USA). Protein levels were quantified using ImageJ and normalized to the β-actin levels.

### Immunofluorescence analyses

Sagittal sections (30 mm) from transgenic (Thy1-aSyn) mice that had received an intracerebroventricular MGO injection were prepared and processed in ^17^. After blocking (0.1M sodium acetate, 5% BSA, 0.05% tween 20) at room temperature for one hour, primary antibodies (used in a concentration of 1µg/mL and diluted in blocking buffer) were applied over night. After three 10 min washes with PBS, the sections were incubated with secondary antibodies (diluted 1:2000 in blocking buffer) for 1 hour at room temperature. Finally, the sections were washed again three times for 10 minutes, stained with DAPI for 5 minutes and cover slipped with Mowiol. For immunofluorescence of cell cultures, cells were grown on glass coverslips, washed and fixed as described above. Permeabilization was performed with 0.5% tritonX 100 at room temperature for 20 min. Coverslips were blocked with 1.5% bovine serum albumin for two hours. Primary antibodies (Tuj1 – 1:3000, Covance; pS129 - 1:2000, Abcam51253; Syn1 – 1:2000, Syn1 – 1:2000, BD Transduction Laboratories) diluted in 1.5% BSA were incubated overnight at 4°C. The coverslips were washed three times with PBS and incubated for two hours with secondary antibodies: anti-mouse and rabbit Alexa Fluor 568-and Alexa Fluor 488-conjugated (1:2000, Life Technologies-Invitrogen). Finally, cells were stained with DAPI for 5 min and cover slipped with Mowiol. Images were analyzed using LAS AF v.2.2.1 (Leica Microsystems) software.

### Immunostaining of human brain samples

Human *post-mortem* brain tissue was obtained from Newcastle Brain Tissue Resource, a UK Human Tissue Authority-approved brain tissue repository. Formalin-fixed paraffin-embedded tissue was obtained from the medulla oblongata of two PD cases and cut at 5µm sections for immunofluorescent analysis. Sections were dewaxed in Histoclear (National Diagnostics, NC, USA) and rehydrated through graded ethanol solutions until water. Antigen retrieval was performed by boiling sections in citrate buffer pH6 for ten minutes prior to immersion in formic acid for five minutes. Sections were blocked in 10% normal goat serum in TBS-T for one hour at room temperature and then incubated in the following primary antibodies suspended in blocking buffer overnight at 4°C: pS129 (El-Agnaf laboratory, 1µg/ml ^27^) and 5G4 (MABN389, Merck, 2µg/ml) combined with either MGO-aSyn (1µg/ml) or MGO-KTK-aSyn (1µg/ml). Primary antibodies were washed off in TBS-T and sections were then incubated in the following secondary antibodies diluted 1:100 in blocking buffer for one hour at room temperature: (goat anti-rabbit-AF405, A48254, Thermo; goat anti-mouse IgG1-AF647, A21240, Thermo; goat anti-mouse IgG2a-AF546, A21133, Thermo). Following secondary antibody exposure, sections were washed and lipofuscin autofluorescence blocked with 0.3% Sudan Black B solution. Sections were visualized on a Leica SP8DLS confocal microscope.

### Determination of Glyoxalase I activity

The quantitative determination of Glyoxalase I (Glo I) activity in SH-SY5Y and rat primary neuronal cultures was performed using a Glyoxalase I Activity Assay Kit (Colorimetric) (K591-100, BioVision, San Diego, CA, USA) according to the manufacturer’s protocol.

### Generation of synthetic peptides containing CEL or CML

Generation of CEL-and CML modified peptides is described in ^28^. In brief, solid-phase peptide synthesis was combined with orthogonal protection of amino acid side-chain functionalities and reductive amination strategies to generate 4 different peptides: Peptides 1 and 2 with CEL- or CML-modified lysines, respectively.

### Statistical analysis

Data were obtained from at least three independent experiments and are shown as mean values ± standard deviation (SD). Two-group comparisons were performed using Student’s t test, multiple-group comparisons were performed using ANOVA with post hoc Tukey. p < 0.05 was considered statistically significant (* p < 0.05; ** p < 0.001; *** p < 0.0001). Statistical analyses were performed in Excel and R.

## Results

### Ribose and MGO-glycation differently affect aSyn pathology

Despite the many open questions regarding the precise role of aSyn in PD pathology, the formation of inclusions containing insoluble aSyn phosphorylated on serine 129 is an important hallmark of PD and other synucleinopathies. Therefore, we first assessed the effects of selected glycated aSyn species, generated as we previously described ^8^. First, we treated SH-SY5Y cells conditionally expressing aSyn ^25^ with aSyn incubated with or without glycating agent. SH-SY5Y cells were differentiated into neuron-like cells with retinoic acid, and at day 4 of differentiations we exogenously added 100nM glycated aSyn and incubated for additional 4 days. Schematic illustration of the procedure is shown in Supplementary Figure 1A. Double immunostaining for Tuj1, a neuron-specific marker, and total aSyn was performed at day 8 and confirmed the induction of the neuronal marker, indicating the neuronal phenotype, and the formation of aSyn inclusions. Furthermore, co-staining with Tuj1 and p-aSyn on Ser129 revealed the presence of phosphorylated inclusions particularly after treatment with aSyn and aSyn-MGO (Supplementary Figure 1B and C). As the glycation protocol involved gentle shaking at 37°C, possibly causing the formation of fibrils and oligomers, it was expected that these control aSyn preparations would lead to the formation of pS129 positive inclusions (Figure 1A). However, treatment with aSyn-ribose resulted in no obvious increase of pS129-aSyn. Correspondingly, in order to further assess aSyn seeding under the treatment with the three different aSyn proteins, we performed a biochemical assay well described previously ^29^ based on detergent fractionation of soluble and insoluble aSyn. Western blotting of the Triton X100-soluble fraction with Syn-1 antibody shows a significant increase of monomeric aSyn under treatment with the three different aSyn proteins, while aSyn in the SDS-soluble fraction migrated at higher molecular weights (HMW) especially under treatment with aSyn and aSyn-MGO, with a molecular mass ranging from 25 to 250 kDa on SDS-PAGE (Figure 1B and 1C). We did not observe any significant increase in HMW products after treatment with aSyn-ribose.

**Figure 1.**
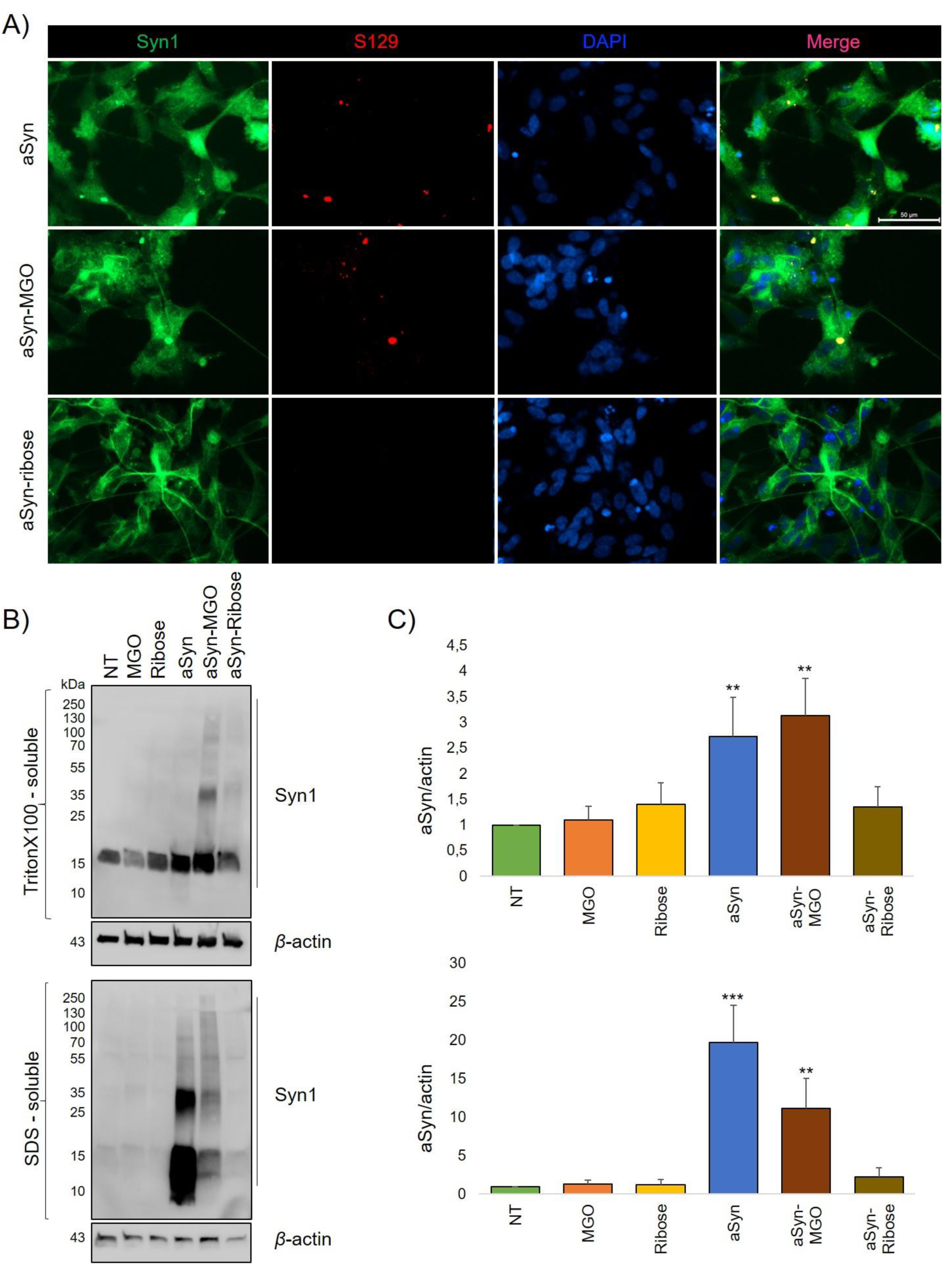
MGO-glycated aSyn induces the formation of pS129 positive inclusions in differentiated SH-SY5Y cells. A). SH-SY5Y cells, conditionally expressing aSyn under a Tet-Off cassette were differentiated to neuron-like cells, expressing human wildtype aSyn (doxycycline was omitted from the medium). aSyn, that had been incubated with or without glycating agent was added to the cells at a final concentration of 100nM for 4 days. Cells were immunostained using Syn1 and pS129, representative images from N=3 (Scale bar 50μm). B). Cells were sequentially extracted with TritonX-100 and SDS containing buffer and analyzed using immunoblots with Syn1 antibodies. β-Actin was used a loading control and C) the ratio of aSyn versus *β*-actin was quantified (n=3, mean ± SD).

We next performed similar experiments in cortical neuronal cultures from wild type rats. At DIV5 we exogenously add the glycated proteins at a concentration of 50nM and further incubated for 20 days. Immunostaining with pS129-aSyn demonstrated the formation of large phosphorylated aSyn inclusions in neuronal cultures incubated with aSyn and aSyn-MGO proteins (Figure 2A). In agreement with the results obtained in SH-SY5Y cells, we detected no phosphorylated aSyn aggregates upon treatment with aSyn-ribose. As a control experiment we treated neuronal cultures with aSyn pre-formed fibrils (PFFs, data not shown). Western blotting of the Triton X100-soluble and SDS insoluble protein fractions was then performed in primary neuronal cultures, revealing significant increase of aSyn HMW species was observed under treatment with aSyn and aSyn-MGO in the SDS-soluble fraction, while aSyn-ribose shows no significant increase (Figure 2B and C). HMW species were absent in non-treated cells (NT) and cells treated with only MGO or ribose. Overall, these data showed a distinct pattern of seeding between aSyn-MGO and aSyn-ribose.

**Figure 2.**
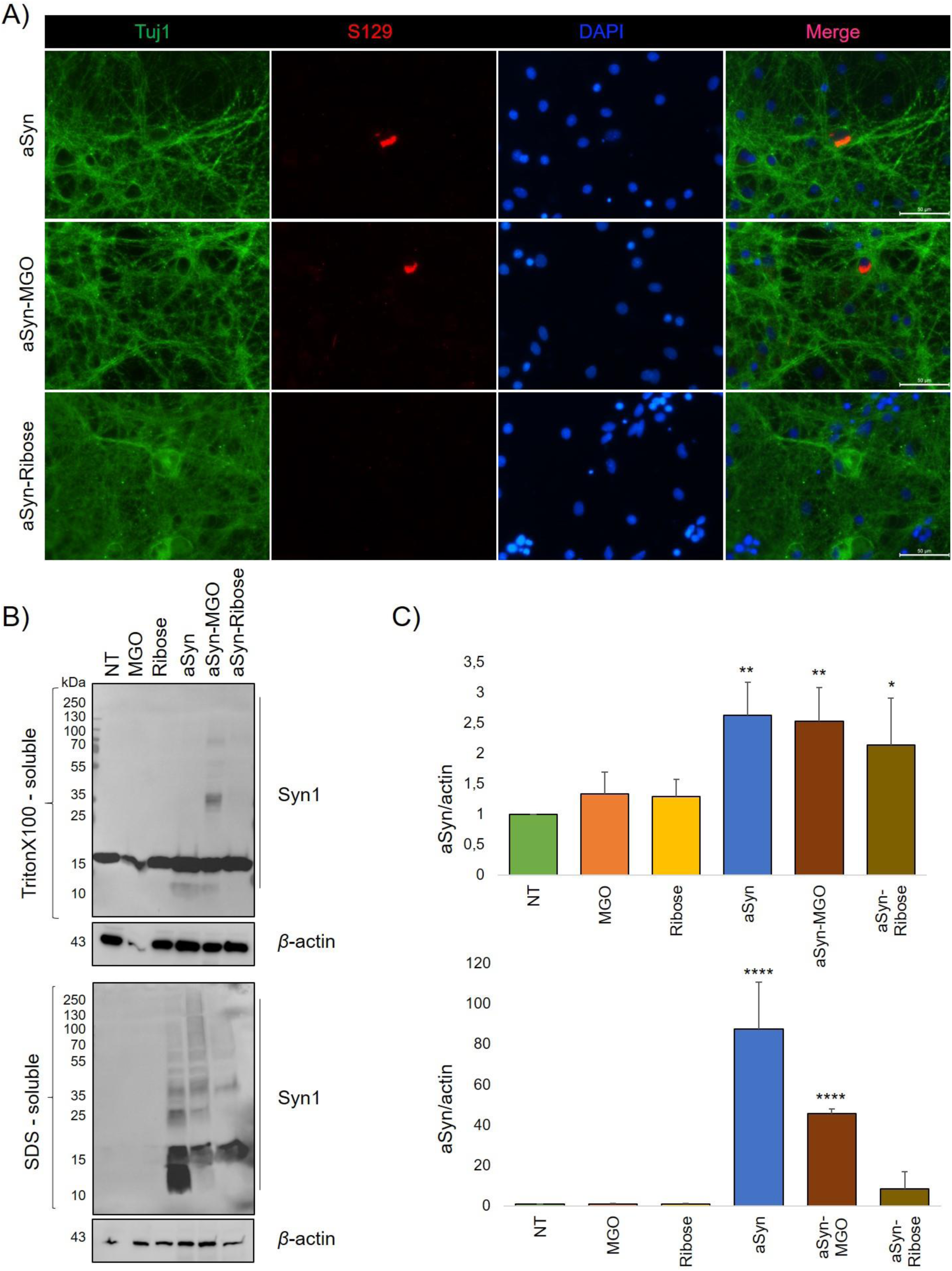
MGO-glycated aSyn induces pS129 positive inclusions in primary neurons. A). Primary rat cortical neurons were incubated for 20 days with 50nM aSyn species. Following wash and fixation, cells were immunostained with Tuj1 and pS129, representative images from N=3 (Scale bar 50μm). B). After sequentially extracting the cells with TritonX-100 and SDS containing buffer, extracts were analyzed using immunoblots with Syn1 and β-actin as loading control. C). The ratio of aSyn and β-actin signal was determined (n=4, mean ± SD).

### Ribose and MGO-glycation decrease Glo I activity

Although the formation of AGEs occurs spontaneously, cells have endogenous active mechanisms to deal with glycation. A highly specific mechanism in detoxifying MGO and other reactive aldehydes occurs via the glyoxalase system, which consists of Glyoxalase-I (Glo I), Glyoxalase-II and aldose reductases ^30,31^. Notably, the endogenous anti-glycation agent Glo I has been found to decrease with age in the human brain ^32^, and in the substantia nigra of PD patients ^33^ as well as in several PD models ^34^. Furthermore, aSyn deficiency in mice causes altered AGEs and increased Glo I expression and glycation stress ^35^suggesting a direct link further illustrating the involvement of aSyn in sugar metabolism. Consistently, we found that the MGO cytotoxicity was abolished upon aSyn knockdown in iPSCs. This further substantiated the association between cytotoxicity and aSyn levels, as we previously reported ^10^. In order to determine whether the administration of glycated aSyn proteins to SH-SY5Y cells and primary neurons was associated with disturbed Glo I, we evaluated both the protein expression levels and the enzymatic activity of Glo I. All samples used in the Glo I protein expression experiments were also used in the Glo I enzymatic experiments. Glo I protein expression did not change after administration of aSyn glycated proteins (Fig. 3A), while Glo I enzymatic activity was shown to reduce significantly, particularly after treatment of cells with aSyn-MGO and aSyn-ribose (Fig. 3B), suggesting impaired function of the glyoxalase pathway.

**Figure 3.**
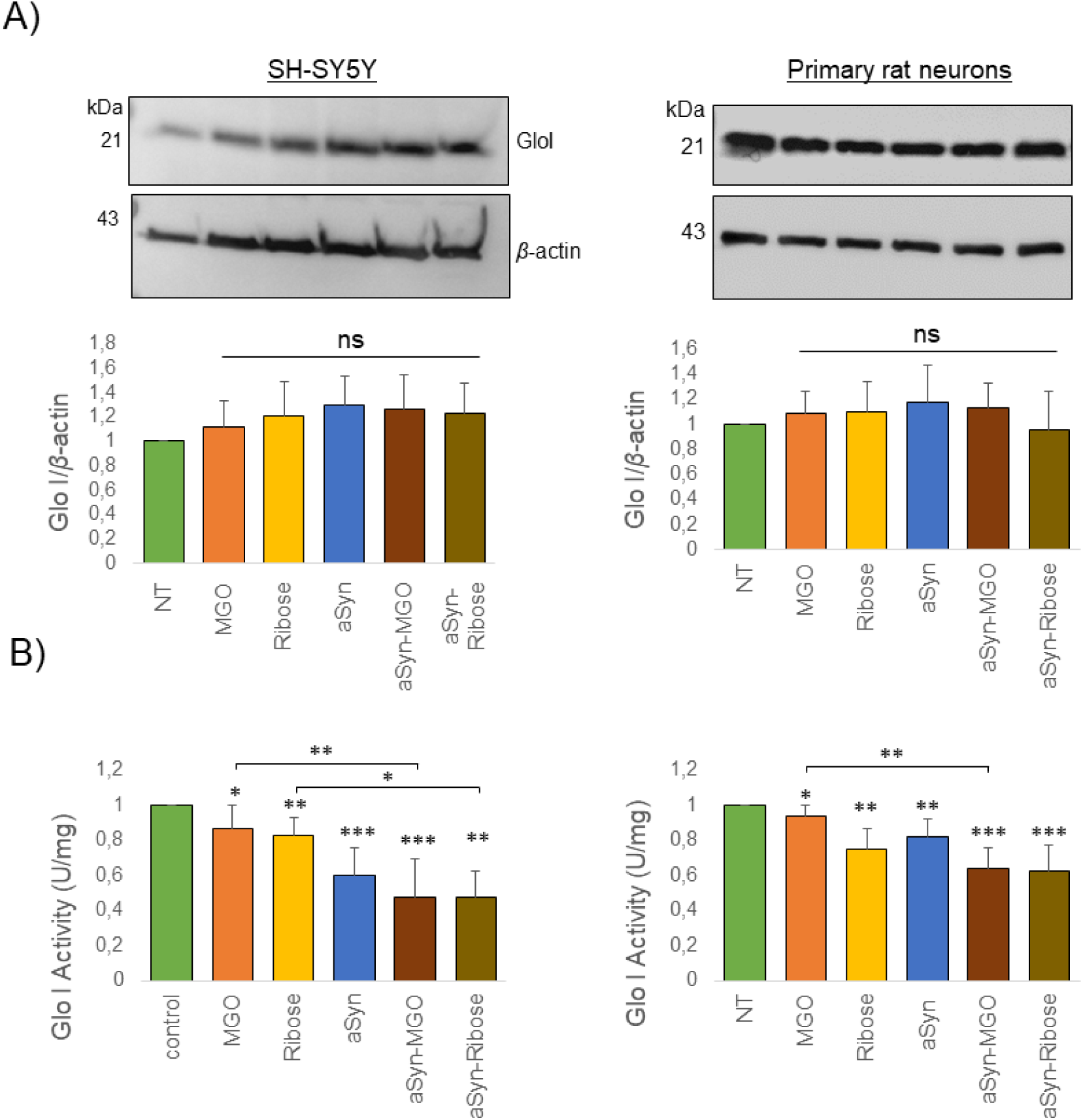
aSyn glycation decrease GloI activity. A). Protein extracts were analyzed using immunoblots with GloI and β-actin as loading control. The ratio of GloI and β-actin signal was determined (n=4, mean ± SD). B). Determination of Glo I activity in SH-SY5Y (i) and primary neuronal cultures (ii) cell lysates. All measurements were performed following kit protocol (n=5, mean ± SD).

### Glycated aSyn induces neuroinflammatory responses in microglial cells

Chronic microglia activation and upregulation of proinflammatory factors modifies different steps of neurodegenerative processes, creating a vicious circle in which neurodegeneration and neuroinflammation fuel each other ^36^. In addition, disrupted function of the glyoxalase pathway can lead to an inflammatory environment contributing to the pathogenesis of neurodegenerative disease ^37,38^. Inflammatory molecules including IL-1β, IL-2, IL-6, IL-10, TNF-α, are often used as markers of neuroinflammation, and are monitored as markers of disease progression, and treatment response in PD. Here, we used ionized calcium-binding adapter molecule 1 (Iba1), a protein predominantly found in cells of the myeloid lineage such as microglia, to label microglia (Figure 4A). Treatment with glycated aSyn species lead to a significant increase in the production of IL6, TNF-α and IL-1β (Figure 4B). Consistently, inducible Nitric Oxide Synthase (iNOS), an enzyme that is induced in response to inflammatory stimuli, and p62, a multi-functional adaptor protein involved in several cellular processes, were increased upon incubation with ribose-or MGO-glycated aSyn (Figure 4C, D and E). Interestingly, both ribose-and MGO-glycated aSyn induced neuroinflammatory responses, but had different effects in the induction of pS129 aSyn accumulation.

**Figure 4.**
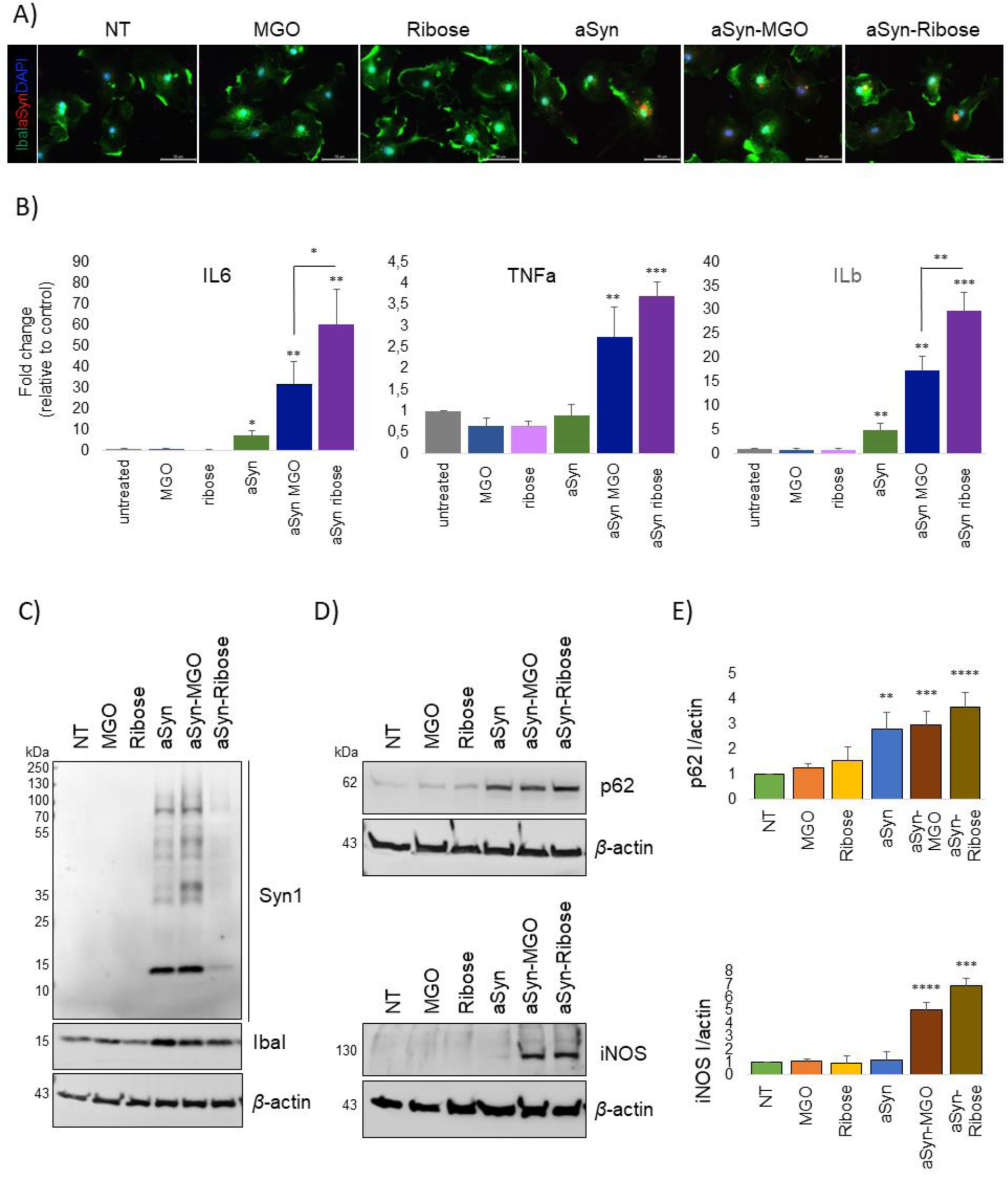
aSyn glycation induces neuroinflammatory responses. A. Microglia were treated with 100nM aSyn species for 24 hours. Cells were stained with Iba1 and Syn1 specific antibodies. Representative images from n=3 (Scale bar 50μm). B. Real time qPCR assessment of the inflammatory target genes IL6, TNFa and ILb in microglia treated cells. β-Actin is used for normalization. C). D). RIPA protein extracts of microglia extracts were examined with immunoblotting for Syn1, Iba1, p62 and iNOS. β-actin is used as a loading control and the intensity of p62 and iNOS signal was quantified (n=4, mean ± SD).

### Novel polyclonal antibodies detect glycated aSyn

We previously demonstrated that MGO-glycation affects primarily *N*-terminal lysine residues in aSyn (K6, K10, K12, K21, K23, K32, K34, K43 and K45) ^10^. Therefore, we next attempted to generate antibodies against MGO-glycated aSyn. We generated three different antibodies: one against glycated full-length aSyn, and two against two different glycated *N*-terminal peptides. Peptides 1 and 2 were designed to span all *N*-terminal lysines (Figure 5A).

**Figure 5.**
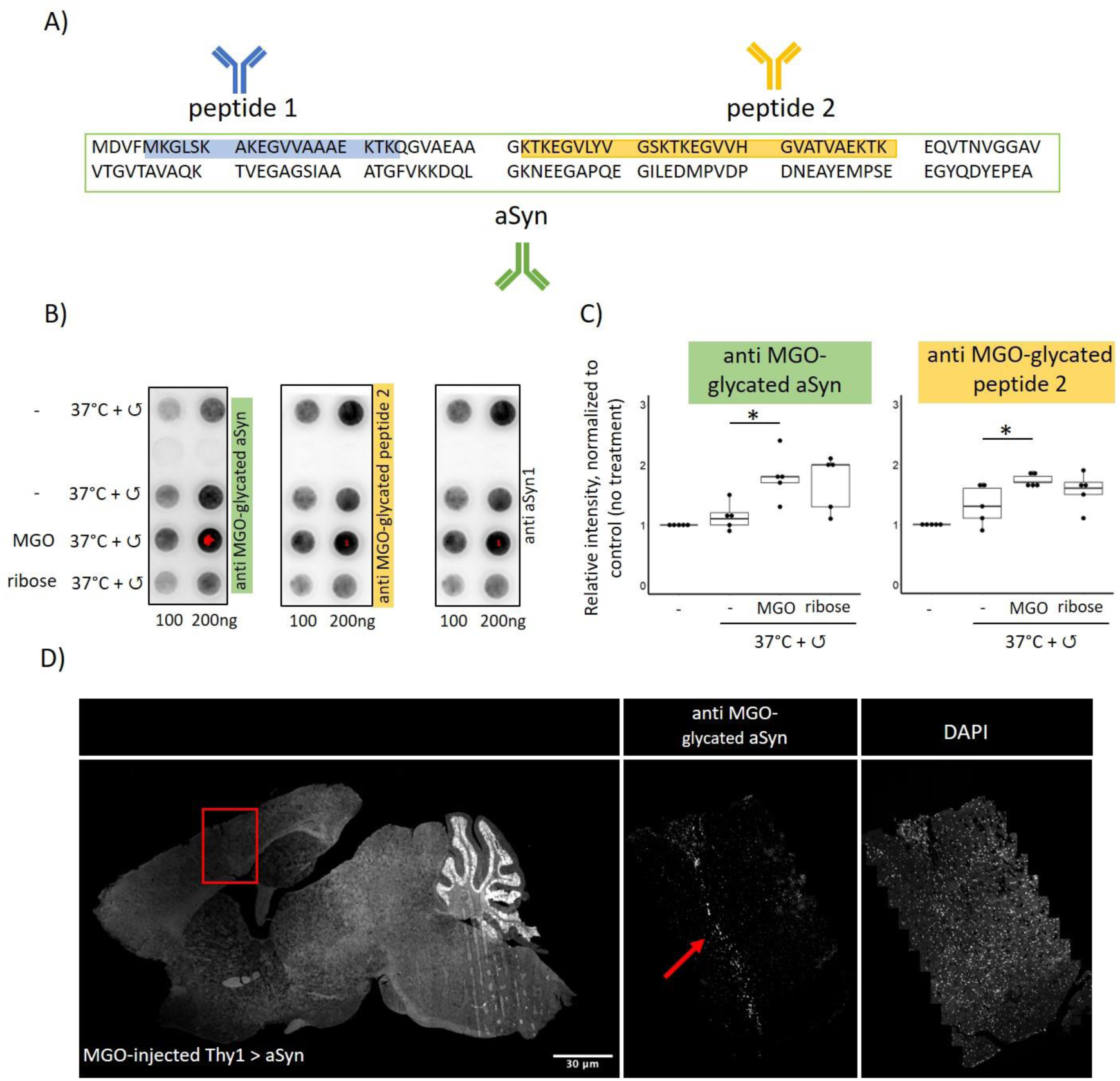
Polyclonal antibodies detect glycated aSyn. A. Three polyclonal rabbit antibodies were generated: Against MGO-glycated full-length aSyn (anti MGO-glycated aSyn) and against MGO-glycated peptides 1 and 2 (anti MGO-glycated peptide 1 or 2). B. Recombinant aSyn was glycated using MGO or ribose. 100 or 200 ng of protein, spotted on a nitrocellulose membrane was stained using the newly produced recombinant antibodies. aSyn1 was used as loading control. One representative blot is shown in B, the quantification of all blots is shown in C). D. Transgenic mice, overexpressing aSyn with a Thy1 promoter injected with MGO. Sagittal sections were stained with anti-MGO-glycated aSyn. Mean ± StDev is displayed, n=3.

Rabbits were immunized with the three immunogens, and were then affinity depleted. To assess antibody specificity, we tested them on glycated peptides (Supplementary Figure 2A – C). As expected, MGO-glycated peptides showed a strong signal, compared to non-glycated control. This was also the case when full-length recombinant aSyn was used (Figure 2C). As ribose and MGO-mediated glycation have distinct effects on aSyn, we also tested whether the antibodies would detect ribose-glycated peptides (Supplementary Figure 2A – C, Figure 2C). Overall, ribose mediated glycation was recognized to a lesser extent and with greater variability, possibly reflecting differences in the glycation pattern or species formed.

Carboxyethyl lysine (CEL) and carboxy methyl lysine (CML) are relevant glycation-induced lysine modifications^39–41^. Therefore, we performed chemical synthesis of aSyn peptides to specifically introduce CEL or CML modifications in specific lysine residues ^28^, and tested the various polyclonal antibodies on synthetic CEL-and CML-modified aSyn peptides (Supplementary Figure 3). While peptide 1-CEL and peptide 2-CEL were both recognized by the anti-MGO-glycated aSyn antibody, peptides 1/2 - CML were not (Supplementary Figure 3 A).

### MGO-glycated aSyn accumulates in mouse and in human brain tissue

To assess the reactivity of the antibodies in tissue, we used brain tissue from transgenic mice expressing human aSyn under the Thy1 promoter^42,43^ that received intracerebroventricular injections with MGO ^17^. Sections from brains of animals injected with vehicle or MGO were analysed blindly. Out of 6 animals that had received MGO injection, 5 displayed strong anti MGO-glycated aSyn immunoreactivity (Supplementary Table 1). Consistently, cells along the needle injection tract, which were more directly exposed to MGO, were strongly stained with anti MGO-glycated aSyn (Figure 5D).

Next, we tested the antibodies in human tissue from the dorsal motor nucleus of the vagal nerve of PD cases, a region amongst the first to be affected by Lewy body pathology in PD^44^. Interestingly, the signal with anti-MGO-glycated aSyn and anti-MGO-glycated peptide 2 consistently colocalized with the signal for the aggregated aSyn antibodies 5G4 and pS129, labelling Lewy bodies and neurites, the characteristic hallmarks of PD-associated pathology (Figure 6).

**Figure 6.**
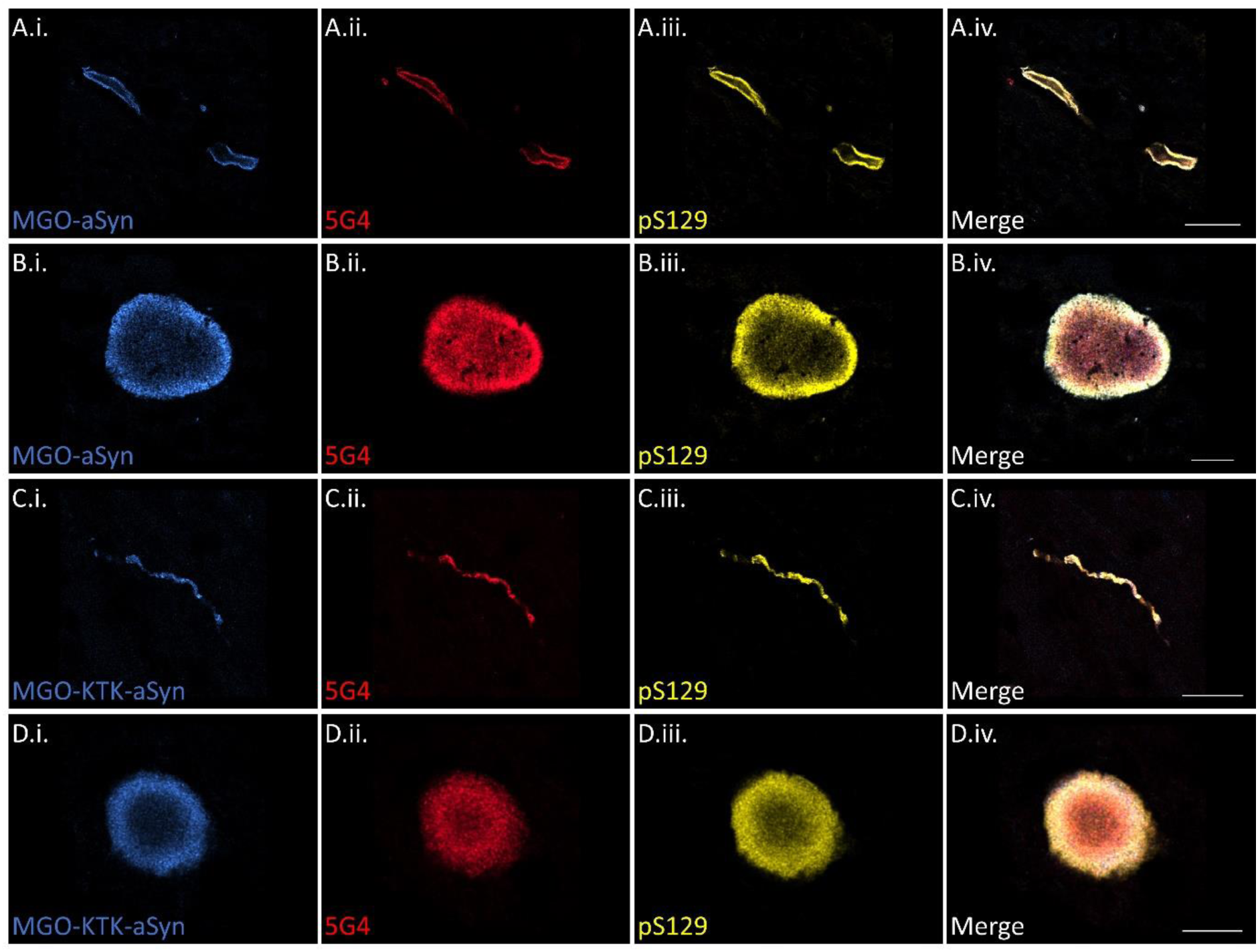
Antibodies against glycated aSyn label Lewy pathology in post-mortem PD brain tissue. Anti-MGO-aSyn labelled both Lewy neurites (A.i.-A.iv.) and Lewy bodies (B.i.-B.iv.) immunoreactive for 5G4 and pS129. Anti-MGO-KTK-aSyn demonstrated a similar pattern of labelling, with Lewy neurites (C.i.-C.iv.) and Lewy bodies (D.i.-D.iv.) labelled by MGO-KTK-aSyn and colocalized with 5G4 and pS129. Scale bars = 30µm (A.i.-A.iv.), 10 µm (B.i.-B.iv. & D.i.-D.iv.), 20 µm (C.i.-C.iv.).

## Discussion

PD and related synucleinopathies are complex, age-associated neurodegenerative disorders of complex etiology. Several environmental factors have been associated with these diseases, but the precise molecular mechanisms involved are still unclear. T2DM, and other metabolic conditions characterized by altered sugar homeostasis are characterized by increased inflammatory responses ^45,46^ and are possible risk factors for the onset and progression of PD ^21,47,48^. Here, we investigated the effect of glycated aSyn species in both neuronal and microglial cells. We found that ribose-and MGO-glycated aSyn have distinct effects on aSyn-associated pathology, with MGO-glycated aSyn inducing significant pS129 inclusions and high molecular weight aSyn species, unlike ribose-glycated aSyn. Both types of glycation induce neuroinflammatory responses, leading to increased production of pro-inflammatory markers such as IL-6, TNF-α, and IL-1β in microglial cells and affect the MGO-detoxifying system.

Interestingly, ribose and MGO-glycated aSyn have different effects on seeding and aggregation of aSyn species in cell models. Based on our findings, it is reasonable to assume that ribose and MGO lead to different kinds or combinations of AGEs, since these are a very diverse family of at least 20 different compounds ^49^. In a previous study, we showed that ribose and MGO affect aSyn aggregation and conformation in different ways ^8^. Consistently, it is well established that different mutations and post-translational modifications cause broad changes in aSyn conformation and pathology^50,51^. Neuroinflammation has been associated with the onset and progression of neurodegenerative diseases ^52,53^ and both innate and adaptive immune systems have been implicated in these diseases. Strikingly, anti-inflammatory medications have shown neuroprotective effects in PD models ^54^. Furthermore, the receptor for advanced glycation end products (RAGE), a type-1 transmembrane glycoprotein of the IgG superfamily, is associated with chronic inflammation in neurodegenerative diseases and is thought to be a potential modulator of neuroinflammation in PD ^55^. Microglia, the resident macrophages of the central nervous system, are key players of the innate immune system. They are critical for regulating neuroinflammation and maintaining homeostasis. Upon pathological stimuli, they undergo physiological changes and become active or “reactive”. Reactive microglia produce high levels of pro-inflammatory cytokines and chemokines that contribute to disruptions of the blood brain barrier, amongst other effects such as altering cerebral cellular homeostasis and neuronal and glial viability. aSyn misfolding causes chronic neuroinflammatory responses ^56^ and in the present study we were able to show that both ribose-and MGO-mediated glycation enhance the effects of misfolded aSyn on microglia activation. Intriguingly, glycation of aSyn seems to affect various phenotypes implicated in PD-associated pathology, including neuroinflammation, inclusion formation, and the glyoxalase system. Neurons and glial cells interact in a complex manner, and even slight changes in their interactions can be problematic and, ultimately, contribute to neurodegenerative processes.

While a causative connection between diabetes and PD was a matter of debate for several years, consensus has been reached now that an existing diabetes increases the odds ratio to also develop PD ^21,47,48^. Moreover, a growing body of literature suggests that, compared to constantly high glucose levels, high fluctuations particularly are more deleterious ^57–60^. While swings of glucose levels are a regular physiological phenomenon, glycemic variability is increased in prediabetic and diabetic individuals ^61^. Furthermore, even individuals not classified as diabetic or prediabetic were shown to experience peaks of glucose levels in the prediabetic and diabetic range ^62,63^. Given that high glucose levels lead to increased MGO levels, it is reasonable to assume that high glycemic variability also causes increased MGO-mediated glycation ^64–66^. But how can this be measured in cell and animal models? Measuring MGO levels is possible but difficult, and the consequences of aSyn glycation have been described^67^. However, directly measuring the glycation status of aSyn has been challenging. Many studies have used antibodies generated against glycated lysine residues in BSA ^17,68–70^, but they are not aSyn-specific. Other antibodies were developed to specifically detect certain aSyn conformations ^71–73^. However, antibodies directed against glycated aSyn have not been previously described, and these are important tools for studying the interplay between diabetes and PD. This is particularly relevant given the recent report that the glucagon-like peptide-1 (GLP-1) receptor agonist lixisenatide, used for the treatment of diabetes, has shown promising results in a clinical trial in people with early PD ^74^. This illustrates that the connection between diabetes and PD is not only crucial for understanding the onset of PD but also that understanding of the molecular interactions can aid in treatment strategies.

In conclusion, our findings demonstrate that MGO-glycated aSyn significantly promotes pathological features associated with PD, including phosphorylated inclusions and HMW aSyn species. Both forms of glycation (MGO and ribose) elicit strong neuroinflammatory responses and impair the glyoxalase detoxification pathway, underscoring a possible critical role of glycation in PD progression. Alongside with the detection of glycated aSyn species in LBs, these insights highlight the potential for targeted therapeutic strategies that address glycation and inflammation, and the development of specific antibodies against glycated aSyn offers promising tools for further investigation and treatment of PD.

## Acknowledgements

This study was supported by the Michael J Fox Foundation, the EU Joint Programme – Neurodegenerative Disease Research project OligoFIT, the Deutsche Forschungsgemeinschaft (DFG, German Research Foundation) under Germany’s Excellence Strategy, EXC 2067/1-390729940, and by the SFB 1286 (project B8). DE is funded by an Alzheimer’s Research UK Senior Fellowship (ARUK-SRF2022A-006).

---

**Supplementary Figure 1.**
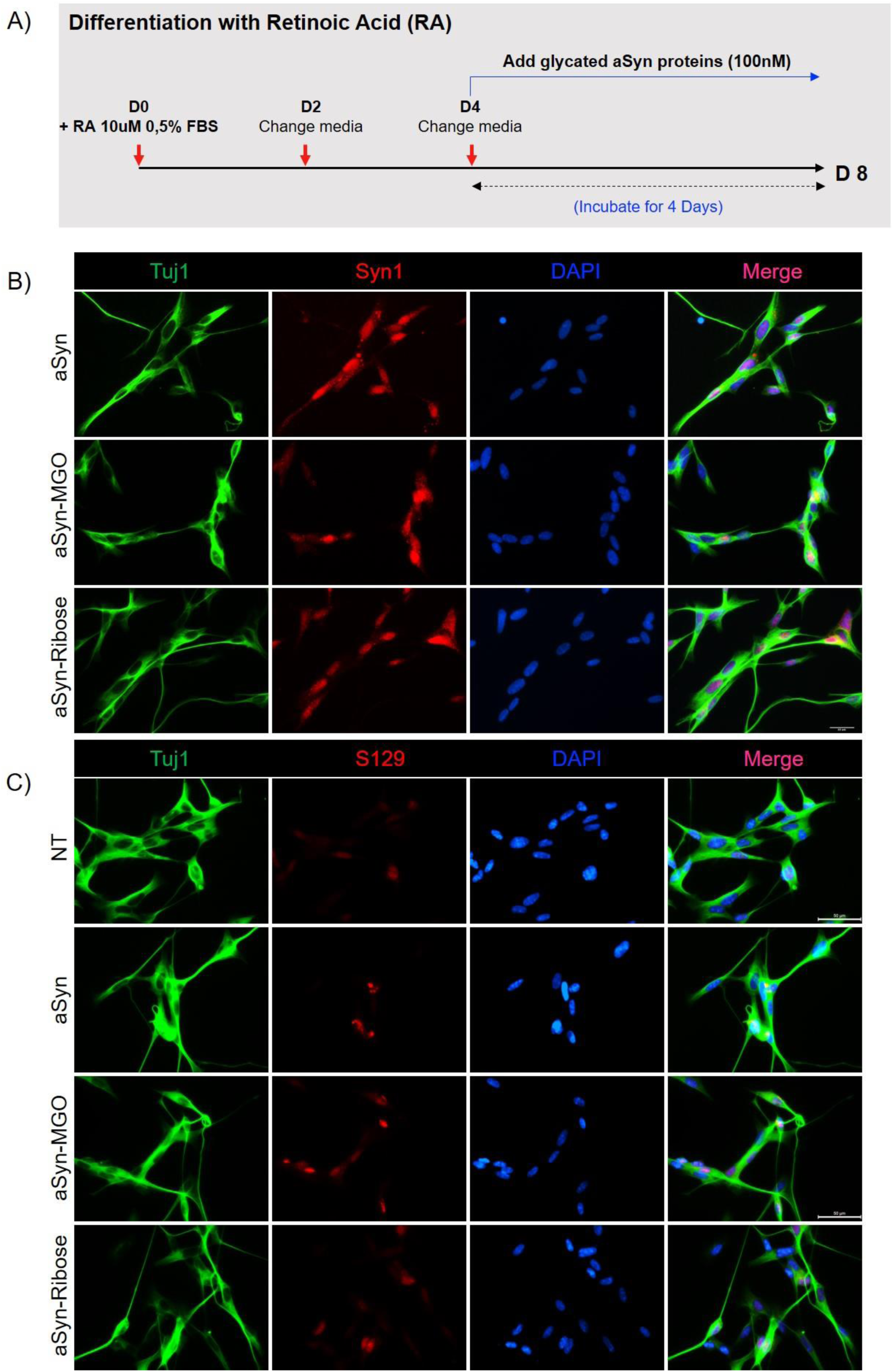
SH-SY5Y cells, conditionally expressing aSyn can be differentiated to neuron-like cells. A). Schematic illustration of the followed differentiation protocol of the SH-SY5Y cells (doxycycline was omitted from the medium). SH-SY5Y cells, conditionally expressing aSyn were differentiated to neuron-like cells, expressing human wildtype aSyn. aSyn, that had been incubated with or without glycating agent was added to the cells at a final concentration of 100nM for 4 days. B). C). Cells were immunostained using Tuj1 and Syn1 (B) or Tuj1 and pS129 (C). Representative images from n=3 (Scale bar 50μm).

**Supplementary Figure 2.**
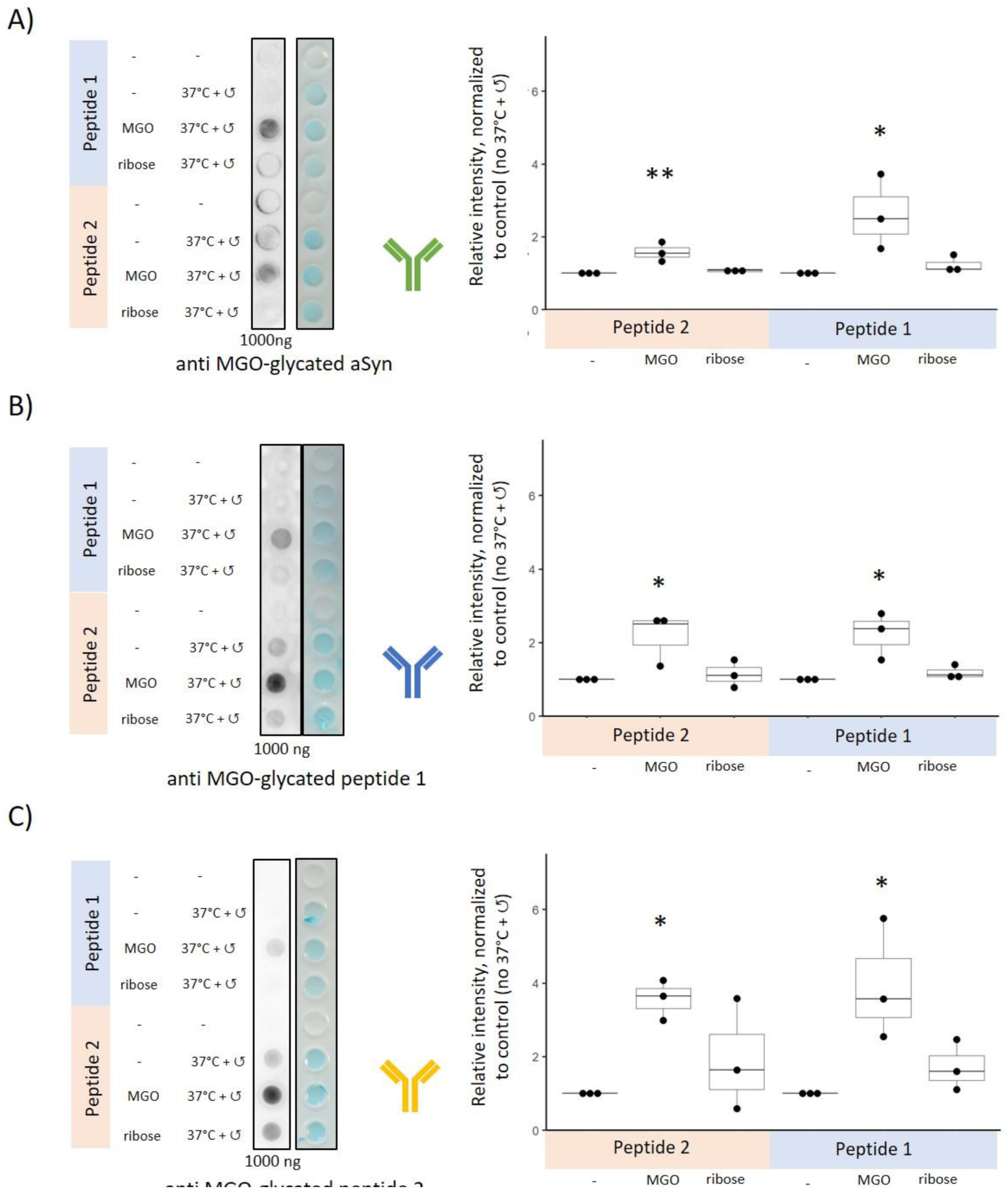
Synthetic peptides, glycated with MGO or ribose are detected by the polyclonal antibodies generated. The antibodies anti MGO-glycated aSyn (A), anti MGO-glycated peptide 1 (B) and anti MGO-glycated peptide 2 (C) were used to stain MGO or ribose glycated peptides. The peptides are depicted in blue (peptide 1) and red (peptide 2). The glycation protocol involves a 5 day incubation with mild agitation at 37°C. Signals were normalized to peptides that had been incubated without glycating agent. Since the peptides cannot be detected with most aSyn specific antibodies, a highly sensitive commercial staining kit was used as loading control. Mean ± StDev is displayed, n=3.

**Supplementary Figure 3.**
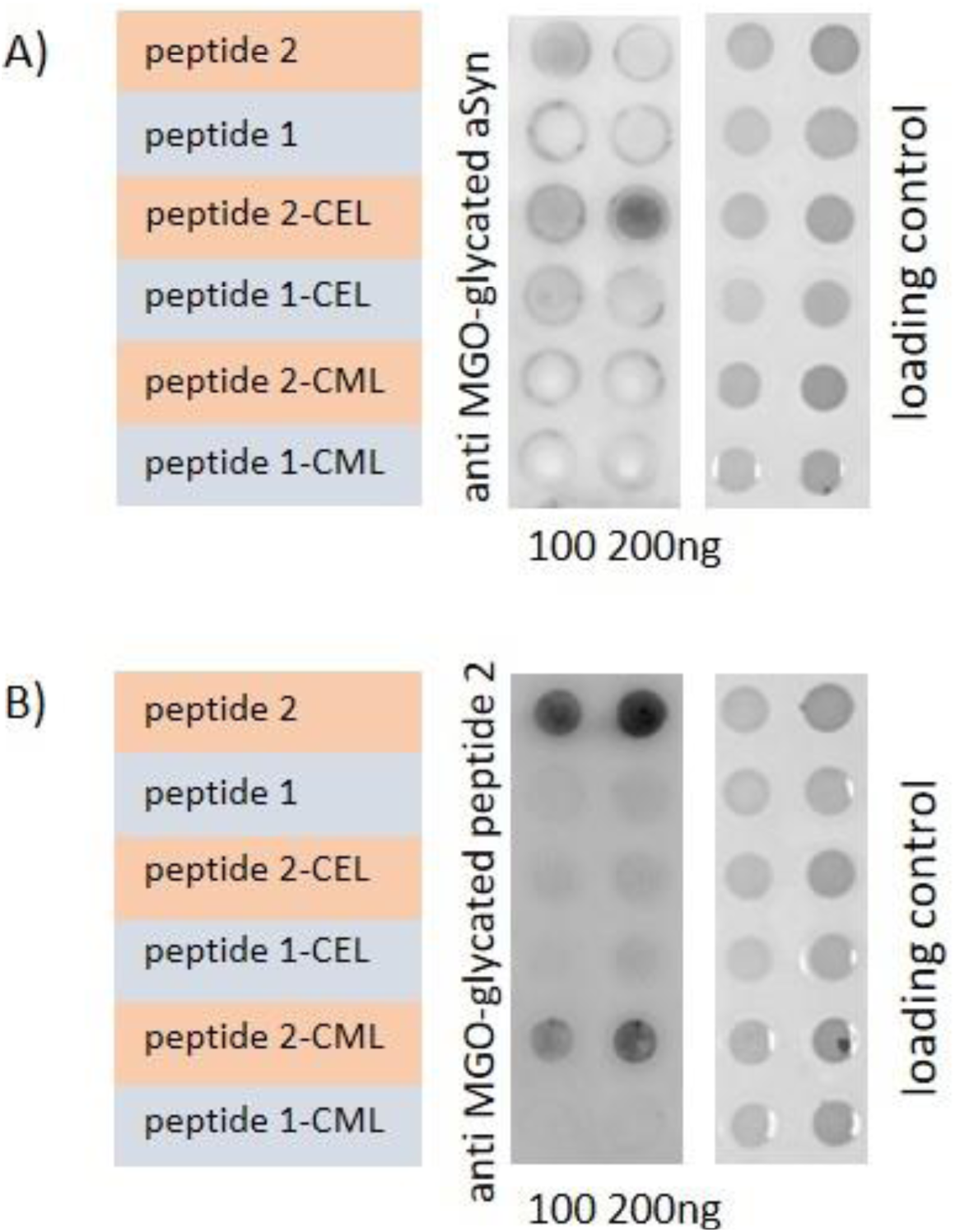
Synthetic AGEs are recognized by the polyclonal antibodies. Synthetic peptides containing CEL-lysine or CML-lysine residues (abbreviated as peptide 1-CEL/CML or peptide 2-CEL/CML) were spotted on nitrocellulose membranes. Anti-MGO-glycated aSyn (A) or anti MGO-glycated peptide 2 (B) were used for staining. The Pierce™ Reversible Protein Stain Kit was used as loading control.

**Supplementary Table 1.** The novel polyclonal antibodies generated detect MGO-glycated aSyn in mouse brain sections. Transgenic animals and control littermates received intracerebroventricular injections of MGO or vehicle ^17^. Sagittal sections of 12 animals were stained with anti-MGO-glycated aSyn and analysed blindly. Blue shading indicates anti MGO-glycated aSyn staining with staining along the ventricle and along the injection channel. Importantly, animal V10 was classified as an outlier due to unusually-low cell numbers, and was excluded.

|  | vehicle |  |  | MGO |  |  |
| --- | --- | --- | --- | --- | --- | --- |
| WT | P5 | P2 | P4 | A5 | A2 | A4 |
| Thy1<br>aSyn | V7 | V8 |  | E9 | E5 | E6 |

